# *Bicc1* and *dicer* regulate left-right patterning through post-transcriptional control of the Nodal-inhibitor *dand5*

**DOI:** 10.1101/2020.01.29.924456

**Authors:** Markus Maerker, Maike Getwan, Megan E. Dowdle, José L. Pelliccia, Jason C. McSheene, Valeria Yartseva, Katsura Minegishi, Philipp Vick, Antonio J. Giraldez, Hiroshi Hamada, Rebecca D. Burdine, Michael D. Sheets, Axel Schweickert, Martin Blum

## Abstract

Rotating cilia at the vertebrate left-right organizer (LRO) generate an asymmetric leftward flow, which is sensed by cells at the left LRO margin. How the flow signal is processed and relayed to the laterality-determining Nodal cascade in the left lateral plate mesoderm (LPM) is largely unknown. We previously showed that flow down-regulates mRNA expression of the Nodal inhibitor Dand5 in left sensory cells. De-repression of the co-expressed Nodal drives LPM Nodal cascade induction. Here, we identify the mechanism of *dand5* downregulation, finding that its posttranscriptional repression is a central process in symmetry breaking. Specifically, the RNA binding protein Bicc1 interacts with a proximal element in the 3’-UTR of *dand5* to repress translation in a *dicer1*-dependent manner. The *bicc1*/*dicer1* module acts downstream of flow, as LRO ciliation was not affected upon its loss. Loss of *bicc1* or *dicer1* was rescued by parallel knockdown of *dand5*, placing both genes in the process of flow sensing.

## INTRODUCTION

Organ asymmetries are found throughout the animal kingdom, referring to asymmetric positioning, asymmetric morphology or both, as for example the vertebrate heart ^1^. The evolutionary origin of organ asymmetries may have been a longer than body length gut, that allows efficient retrieval of nutrients, and the need to stow a long gut in the body cavity in an orderly manner ^2^. Vertebrate organ asymmetries (*situs solitus*) are quite sophisticated: in humans, the apex of the asymmetrically built heart, with two atria and ventricles each that connect to lung and body circulation, points to the left; the lung in turn, due to space restrictions, has fewer lobes on the left than on the right side (2 and 3 in humans), stomach and spleen are found on the left, the liver on the right, and small and large intestine coil in a chiral manner. In very rare cases (1:10.000), the organ situs is inverted (*situs inversus*).

Heterotaxia describes another rare situation (about 1:1.000), in which subsets of organs show normal or aberrant positioning and/or morphology, cases inevitably associated with severe disease syndromes ^3–6^.

The Nodal signaling cascade takes center stage in setting up organ situs during embryonic development ^1, 7^. *Nodal* is activated in the left lateral plate mesoderm (LPM) of the early neurula embryo, where it induces its own transcription, that of its feedback inhibitor *Lefty* and the homeobox transcription factor *Pitx2*. *Pitx2* controls the asymmetric placement and morphogenesis of organs during subsequent development, long after Lefty has terminated Nodal activity during neurula stages ^7^. In deuterostomes, i.e. echinoderms and chordates, cilia are required for Nodal cascade induction ^1, 8, 9^. The archenteron, that is the primitive gut or remnants thereof, transiently harbors the ciliated epithelium of the left-right (LR) organizer (LRO) during neurula stages. The LRO is typically characterized by motile cilia at its center and immotile, supposedly sensory cilia at its lateral borders. The posterior orientation and tilt of motile cilia, together with their intrinsic clockwise rotation, give rise to a leftward fluid flow in the extracellular space. Flow is sensed by the immotile cilia on lateral LRO cells at the left LRO margin. Subsequently, the Nodal cascade is activated at a distance in the left LPM. A great many cilia mutants and experimental manipulations of motile cilia in diverse vertebrate model organisms have underscored this general mechanism in fish, amphibian and mammalian embryos (including humans), while cilia were lost in sauropsids (reptiles and birds) ^1^,^10^,^11^.

The decisive molecular consequence of leftward flow is the repression of the Nodal inhibitor *dand5* at the left LRO margin, as visualized by reduced mRNA expression at post-flow stages ^12–14^. Importantly, by manipulating flow and *dand5*, the Nodal cascade can be modulated at will: morpholino oligomer (MO) mediated gene knockdown on the right side induces the cascade bilaterally while left-sided knockdown rescues the cascade in the absence of flow (for example through impairment of ciliary motility; ^6, 14–16^. All these experimental manipulations are highly efficient (close to one hundred percent), confirming the central role of *dand5* repression by flow ^17^. In due course, the Nodal signal transfers from the LRO to the left LPM, where it induces the left-asymmetric Nodal signaling cascade. In the frog *Xenopus*, the flow-dependent decay of *dand5* mRNA on the left LRO margin occurs somewhat too late, i.e. at the very stage (st. 19/20) in which *Nodal* becomes activated in the left LPM for the first time. In addition, left-sided *dand5* mRNA repression is observed in a maximum of about 80% of WT specimens, while the arrangement of inner organs is disturbed in less than 5% of cases, i.e. the observed flow-dependent down-regulation of *dand5* mRNA does not suffice to explain the robust occurrence of *situs solitus*. Posttranscriptional mechanisms controlling *dand5* asymmetry control may therefore be at work as well.

Elucidating the mechanisms of flow-mediated *dand5* repression has been challenging: targeted events need to be separated from earlier steps, particularly the morphogenesis of the LRO with its arrangement of central motile and lateral immotile cilia. When these upstream events are impaired– either experimentally or genetically – laterality defects inevitably arise, such as for example in cilia motility mutants, which affect the same readouts available for flow-sensing mechanisms. The following criteria apply to factors involved in flow sensing and repression: (a) factors have to act at the lateral LRO in the population of flow sensing cells; (b) upstream events, particularly flow, have to proceed normally when candidate factors are down-regulated in the sensing cell population; (c) flow sensor cells should be present upon factor loss-of-function; and (d) factor loss should be rescued by loss of *dand5*, i.e. by artificially over-riding flow-mediated repression. Only two such factors have been described to date: the cation channel TRPP2 (encoded by *Pkd2*), which is a critical determinant of kidney development and function, and which we initially characterized as an LR determinant in a *Pkd2* knockout mouse ^18^. Pkd1l1, which is expressed in LROs, binds to and co-localizes with TRPP2/*Pkd2*, mutants of which have normal cilia and flow but abnormal *dand5* and which acts genetically downstream of flow and upstream of *Pkd2* ^19–21^.

Here, we identify two additional such factors: the RNA-binding protein Bicaudal-C (Bicc1 in *Xenopus*) and Dicer1, the enzyme catalyzing the final step of microRNA (miR) biosynthesis. Both are instrumental for repression of *dand5*, which we show to work at the level of translational control. In addition, our data indicate that *bicc1*, *dicer1* and *pkd2* interact in *dand5* repression.

## RESULTS

### *Bicc1* regulates left-right axis formation downstream of leftward flow

Bicc1 regulates cell fate decisions during embryonic development and is conserved from *Drosophila* to mammals ^22–26^. Bicc1 binds to selected mRNAs and regulates translation post-transcriptionally, in a positive ^27^ or negative context dependent manner ^25, 28^. *bicc1* mutant mice display LR asymmetry, heart, kidney and pancreas defects ^27, 29^, while in *Xenopus* (like in *Drosophila*), maternal Bicc1 in addition regulates anterior-posterior development ^24, 28, 30^. Previous work showed that *bicc1* in frog and mouse was (a) involved in LR axis formation ^29^; (b) *bicc1* and *pkd2* interacted in kidney development ^27^. *bicc1* loss-of-function resulted in mispolarized LRO cilia by impacting on Wnt/planar cell polarity (PCP) signaling. In the kidney, Bicc1 regulated *pkd2* mRNA stability and translation positively by antagonizing miR17 activity. In both contexts, Bicc1 protein localized to P-bodies, cytoplasmic complexes involved in mRNA stability and turnover ^27, 29, 31^. The expression pattern of *bicc1* in the frog LRO revealed a strong enrichment of mRNA transcription in the flow-sensing lateral LRO cells (cf. Figure 4 in ^29^, indicative of a specific function in these cells, which was not addressed at the time because of impaired flow generation in mutants and morphants.

The frog *Xenopus* offers a precise targeting of flow-sensing cells by microinjection of the left or right so-called C2-lineage ^32–34^, while avoiding the central flow-generating part of the LRO (Figure 1A). To knock down *bicc1*, a previously used translation-blocking antisense morpholino oligomer (TBMO) was used as well as a newly designed MO interfering with splicing (splice-blocking MO; SBMO). In both cases, two MOs were used which specifically targeted the S- or L-allele of the allotetraploid frog *Xenopus laevis* ^35, 36^, which are both expressed during embryogenesis and encode identical proteins (cf. gene information on the community website Xenbase). Injecting either MO in isolation did not affect laterality (Figure S1A).

**Figure 1.**
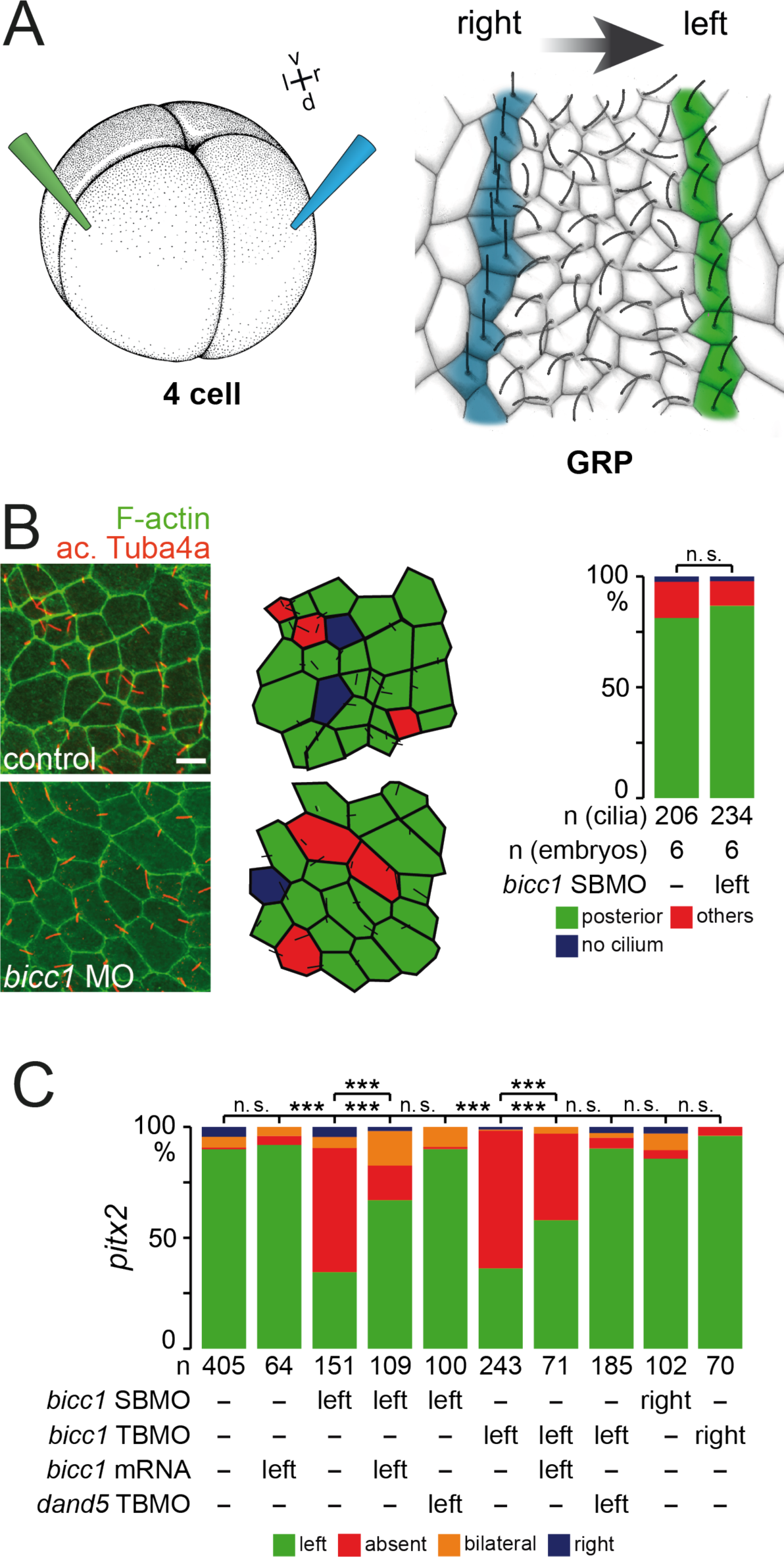
*bicc1* acts downstream of leftward flow and upstream of *dand5* repression.| (A) Schematic depiction of injection scheme at the 4-cell stage (left) to target specifically left (green) or right (blue) flow sensing cells at the gastrocoel roof plate (GRP; right), which is shown as a dorsal explant of the archenteron at stage 19. (B) Unaltered GRP ciliation in *bicc1* morphants. Representative dorsal explants of control and *bicc1* morphant specimens (left) as well as assessment of GRP morphology and cilia polarization (middle and right). (C) Absence of left LPM *pitx2* expression in *bicc1* morphants, unilaterally injected on the left, was rescued by parallel knockdown of *dand5*. In case of TBMO, mouse *bicc1* was co-injected. In case of SBMO, full-length *Xenopus bicc1* was used. Numbers represent analyzed cilia (B) and embryos (C), respectively, from >3 independent experiments each. n.s., not significant; ***, very highly significant, p<0.001. Scale bar in (B) represents 10 µm.

In morphant embryos, in which the left sensor cells were targeted by co-injection of S- and L-MOs, *foxj1* expression was unaltered in the precursor tissue of the LRO, superficial mesoderm (SM; not shown) as was GRP morphology and cilia polarization, demonstrating targeting specificity (Figure 1B). LPM *pitx2* expression, however, was predominately absent in morphants injected unilaterally on the left side (Figure 1C; Figure S1B, C). MO-specificity was demonstrated by (a) right-sided injections, which did not affect *pitx2* (not shown); (b) co-injection of full length *bicc1* mRNA that was not targeted by either MO (mouse *bicc1*, *mbicc1*, in case of TBMO, and *Xenopus bicc1* in case of SBMO), which rescued LPM *pitx2* expression in a significant proportion of specimens (Figure 1C; Figure S1D). *bicc1* gain-of-function alone did not affect *pitx2*, neither on the left (Figure 1C) nor on the right (not shown). Both MOs gave virtually identical results, fulfilling yet another criterion for the controlled use of MOs ^37^. Importantly, parallel knockdown of *bicc1* and *dand5* in left LRO sensor cells rescued *pitx2* expression (Figure 1C), demonstrating that *bicc1* acted downstream of flow and upstream of flow-mediated *dand5* repression, i.e. in the process of flow sensing.

### *Bicc1* represses *dand5* translation

Next, we tested if and how *bicc1* acted on *dand5*. Our previous identification of *dand5* mRNA as a target of Bicc1 binding ^30^ made us wonder whether Bicc1 regulated Dand5, the critical target downstream of leftward flow ^1^. To directly test this possibility, we set up an assay in animal cap explant cultures (AC-assay; Figure 2A) using our previously published *dand5* 3’-UTR luciferase reporter ^30^; Figure 2B). The animal cap of the gastrula embryo at stage 10 expresses maternal transcripts of *dand5* ^38^, while *bicc1* is not present in this tissue ^26^. Injection of reporter constructs harboring the full-length 3’-UTRs of the respective S and L-alleles of *dand5* into the animal region of the 4-cell embryo, therefore, resulted in protein translation and luciferase activity in the AC-assay (Figure S2B). Co-injection of *bicc1* mRNA, however, repressed luciferase activity to around 20%. A full-length mouse *bicc1* construct repressed reporter activities of S- and L-allele as well, though slightly less efficient (Figure S2B). These experiments demonstrated a repressive effect of *bicc1* on a reporter protein expressed from a construct harboring the *dand5* 3’-UTR.

**Figure 2.**
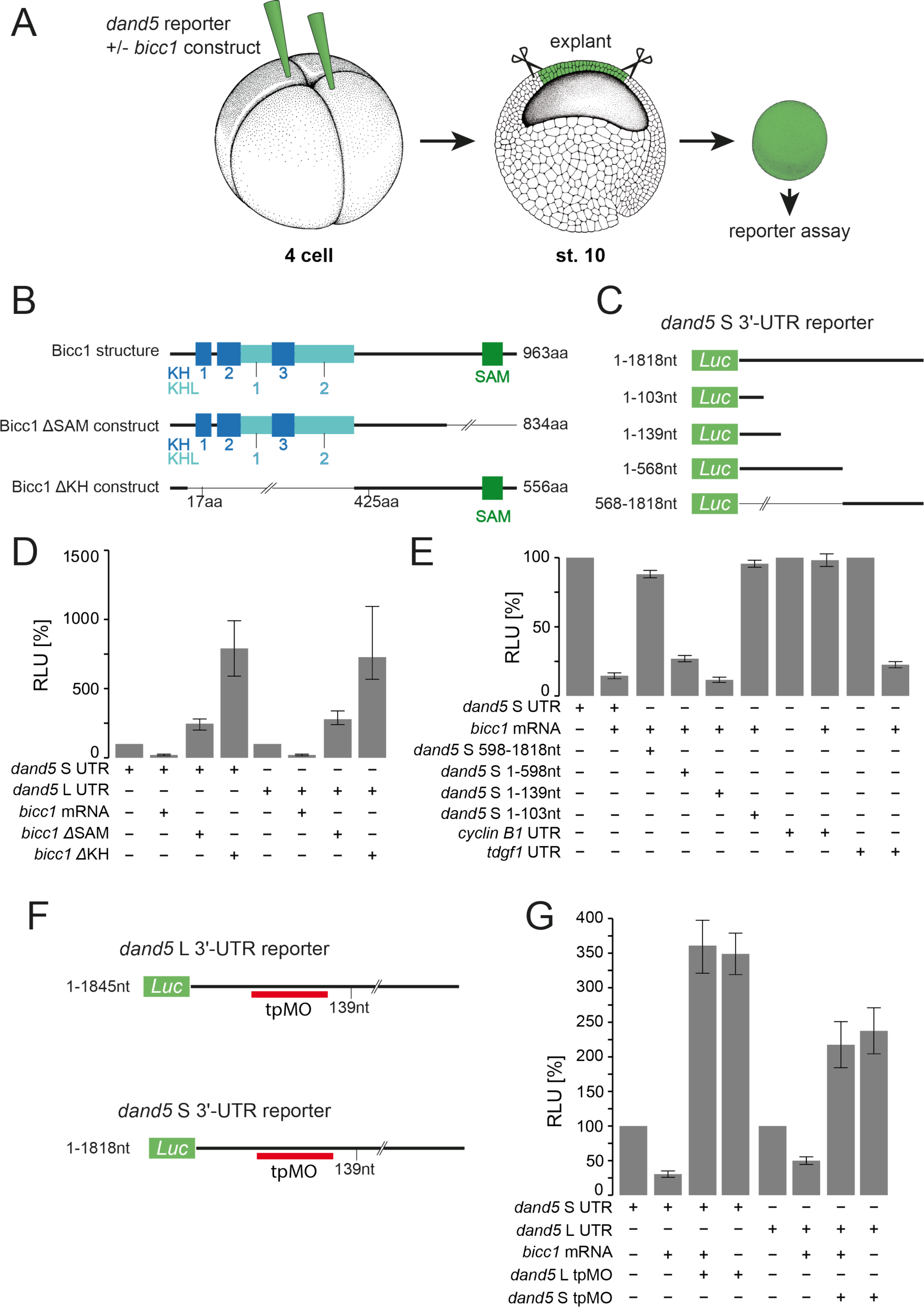
Bicc1 represses *dand5* mRNA translation. (A) Schematic depiction of *dand5* reporter assay. Luciferase reporter constructs fused to *dand5* 3’-UTR sequences were co-injected with *bicc1* effector constructs into the animal region of 4-cell embryos. Following culture to stage 10, the animal cap region was excised and assayed for luciferase activity. (B) Bicc1 protein domains and deletion constructs. (C) Luciferase reporter constructs harboring different regions of the *dand5* (S-allele) 3’-UTR. (D) Bicc1-dependent repression of the *dand5* reporter gene requires both KH- and SAM-domains. (E) Repression of *dand5* translation is mediated through a proximal sequence element in the 3’-UTR. (F) Schematic depiction of TPMO-binding to the *dand5* 3’-UTR of the L-reporter mRNA. (G) Co-injection of TPMO prevented *bicc1*-dependent repression of the full-length *dand5* L-reporter.

Sequence conservation of the 3’-UTRs between the alloalleles is low, except for the proximal 230 nucleotides, which show 84% sequence identity (Figure S2A, C). Flow-dependent mRNA repression of *dand5* was found for both alleles, as visualized by whole-mount in situ hybridization of dorsal explants at stages 18 and 20 with antisense RNA probes specific for the 3’-UTRs of S- and L-alleles (Figure S2D, E). Bicc1 harbors three RNA-binding KH (hnRNP-K homology) and two KH-like domains in its N-terminal part, and an RNA and protein binding SAM (sterile alpha motive) domain at the C-terminus ^26, 28^Figure 2B). We therefore wondered which domain of the Bicc1 protein was required for repression of *dand5* translation. Co-injection of the reporter gene with a deletion construct in which the SAM-domain was missing (ΔSAM; ^26^ not only prevented the repressive action of Bicc1, but resulted in an about 2-3 fold enhanced reporter activity, as compared to injection of the reporter construct alone (Figure 2D). This effect was even more pronounced when a deletion construct was used in which all of the KH- and KH-like domains were absent (ΔKH; ^26^; Figure 2D). In this case, reporter activity peaked at about 7.5 times the value of control levels (Figure 2D), in agreement with the proposed dominant-negative function of this deletion construct ^26^. These results showed that both, KH and SAM domains were required for *dand5* repression.

In the next series of experiments, we asked which sequences in the 3’-UTR were required for translational inhibition by Bicc1. In this part of the analysis, we initially restricted ourselves to the 1818 nucleotides of *dand5*S. Deleting the proximal 598 nucleotides alleviated the repressing effect to almost WT levels (Figure 2E). This proximal sequence alone conferred translational repression to just under 40% of WT, i.e. slightly less than the full-length 3’-UTR. Further deletion to nucleotides 1-139 allowed repression at WT levels, while deleting additional 26 nucleotides (construct 1-103) abolished repression. The specificity of the assay was validated by using a *cyclin B1* reporter, which was not repressed by Bicc1 (negative control), and a *tdgf1* (previously known as *cripto*) reporter, which was repressed (positive control), as previously reported ^30^. To test whether the proximal element of the *dand5* 3’-UTR was instrumental in mediating Bicc1-dependent translational repression, we designed an antisense target protector MO (tpMO) covering nucleotides 91-116 of the L- and nucleotides 107-132 of the S-alloallele, respectively (Figure 2F; Figure S2C). Co-injection of the tpMO with the full-length *dand5* reporter and *bicc1* mRNA prevented the *bicc1*-dependent reporter gene repression (Figure 2G). This result confirmed the role of the proximal 3’-UTR sequences in *bicc1*-dependent *dand5* repression. Remarkably, the reporter activity was enhanced by tpMO about two-fold, as was the reporter upon co-injection with tpMO in the absence of *bicc1* (Figure 2G). This data set indicates that additional component(s) restrict *dand5* activity through interaction with its 3’-UTR independent of Bicc1.

Finally, we wondered whether *bicc1* affected *dand5* translation in left-right development as well. Without a specific antibody that recognized Dand5, we assayed *pitx2* expression, which is a direct readout of *dand5* repression ^17^. Deletion constructs of *bicc1* removing KH- and KHL- or the SAM-domains were unable to rescue *pitx2* expression in *bicc1* morphants (Figure 3A), corroborating the results obtained with these constructs in the AC-reporter assay (Figure 2D). Injection of ΔKH alone resulted in absence of *pitx2* expression in some 30% of specimens, supporting the supposed dominant-negative role of this construct in much the same way as in the reporter assay (cf. Figure 2D; ^26^. Left-sided injection of the tpMO prevented *pitx2* induction in the left LPM in close to 50% of specimens (Figure 3B), demonstrating that this sequence was required for *dand5* repression *in vivo* as well.

**Figure 3.**
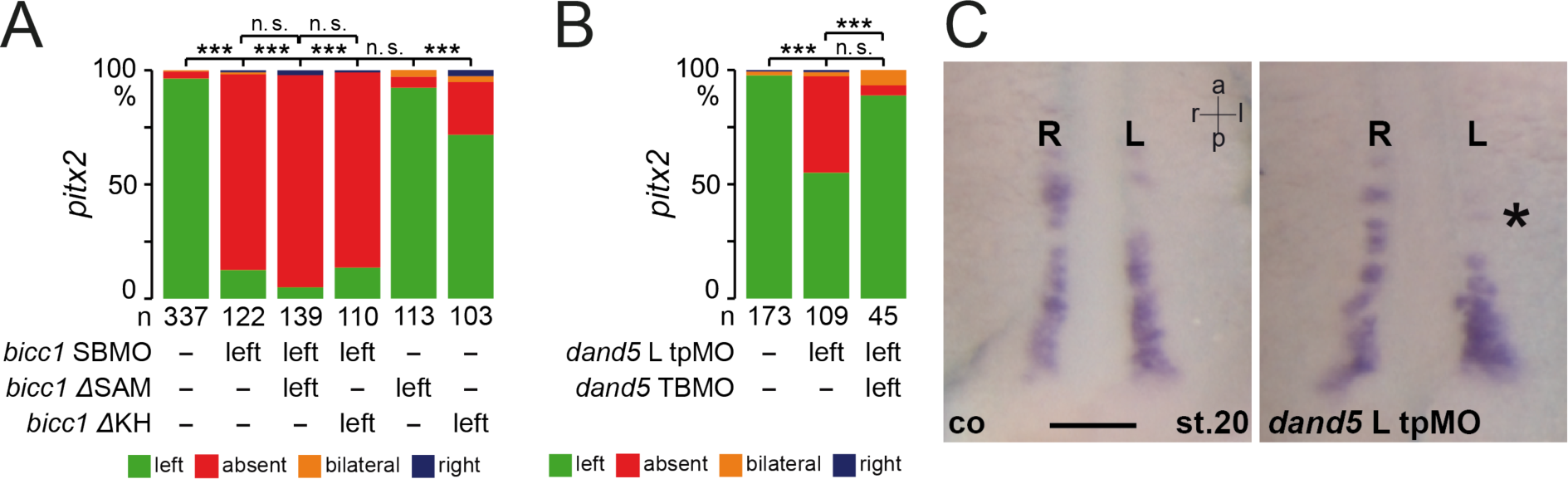
Bicc1 represses *dand5* mRNA translation *in vivo* (A) No rescue of LPM *pitx2* expression in *bicc1* morphants co-injected with either KH/KHL- or SAM-deletion constructs. (B) Injection of a tpMO covering nucleotides 9-116 of the 3’-UTR of *dand5* L-alloallele into the lineage (C2) of left GRP flow sensor cells (cf. Figure 1A) prevented *pitx2* induction in the left LPM. Expression was rescued by co-injecting a *dand5* TBMO. (C) Unaltered mRNA expression of *dand5* in control uninjected (co; left) and tpMO-injected (right) specimens. Numbers in (A, B) represent analyzed specimens from >3 independent experiments. n.s., not significant; ***, very highly significant, p<0.001. Scale bar in (C) represents 100 µm.

Importantly, co-injection of a *dand5* TBMO rescued asymmetric *pitx2* LPM induction. Asymmetric decay of *dand5* mRNA in left flow-sensor cells at the GRP was not impaired by tpMO-injections (Figure 3C). Together, these experiments demonstrated that Bicc1 controlled *dand5* translation in a posttranscriptional manner through interaction with a proximal element in the 3’-UTR and dependent on both KH/KHL- and SAM-domains.

### *dicer* and *bicc1* interact in posttranscriptional *dand5* regulation

The Bicc1-dependent regulation of *pkd2* translation via miRs as well as the localization of Bicc1 to P-bodies indicated that miRs might be involved in the regulation of *dand5* via Bicc1 as well. The RNase III enzyme Dicer processes pre (precursor) -miRs in the cytoplasm and - together with Ago2 - assembles the RNA-induced silencing complex RISC ^39^. In the kidney, Bicc1 acted downstream of Dicer1 to transfer target mRNAs such as *ac6* (*adenylate cyclase 6*) or *PKIalpha* (*protein kinase A inhibitor alpha*) unto Ago2, which cuts or blocks their translation in a miR-dependent manner ^40^. As a first step towards exploring a possible role of miRs in *dand5* regulation, we analyzed the expression of *dicer1*. Zygotic *dicer1* mRNA was expressed in somites and notochord at flow-stage (st. 17) (Figure 4A). Remarkably, mRNA was found specifically in lateral cells of the GRP, excluding the notochordal GRP cells in-between and the lateral endodermal cells flanking the GRP (Figure 4A, A’). Two MOs that targeted translation (TBMO1, TBMO2) through conserved sequences of both S- and L-alleles were used to knockdown *dicer*. Targeting the left side of the GRP (C2 flow-sensor lineage) abolished *pitx2* expression in the left LPM (Figure 4B). The absence of phenotypes upon right-sided MO-injections argues against MO toxicity and off-target effects (Figure 4B). A parallel knockdown of *dand5* on the left rescued WT *pitx2* expression (Figure 4B; Figure S3A-C), further supporting MO-specificity. In mouse embryos, *dicer* was required for Nodal cascade induction as well. Induced conditional deletion of *dicer* from the mouse LRO prevented expression of *Nodal* mRNA in the left LPM (Fig. 4C).

**Figure 4.**
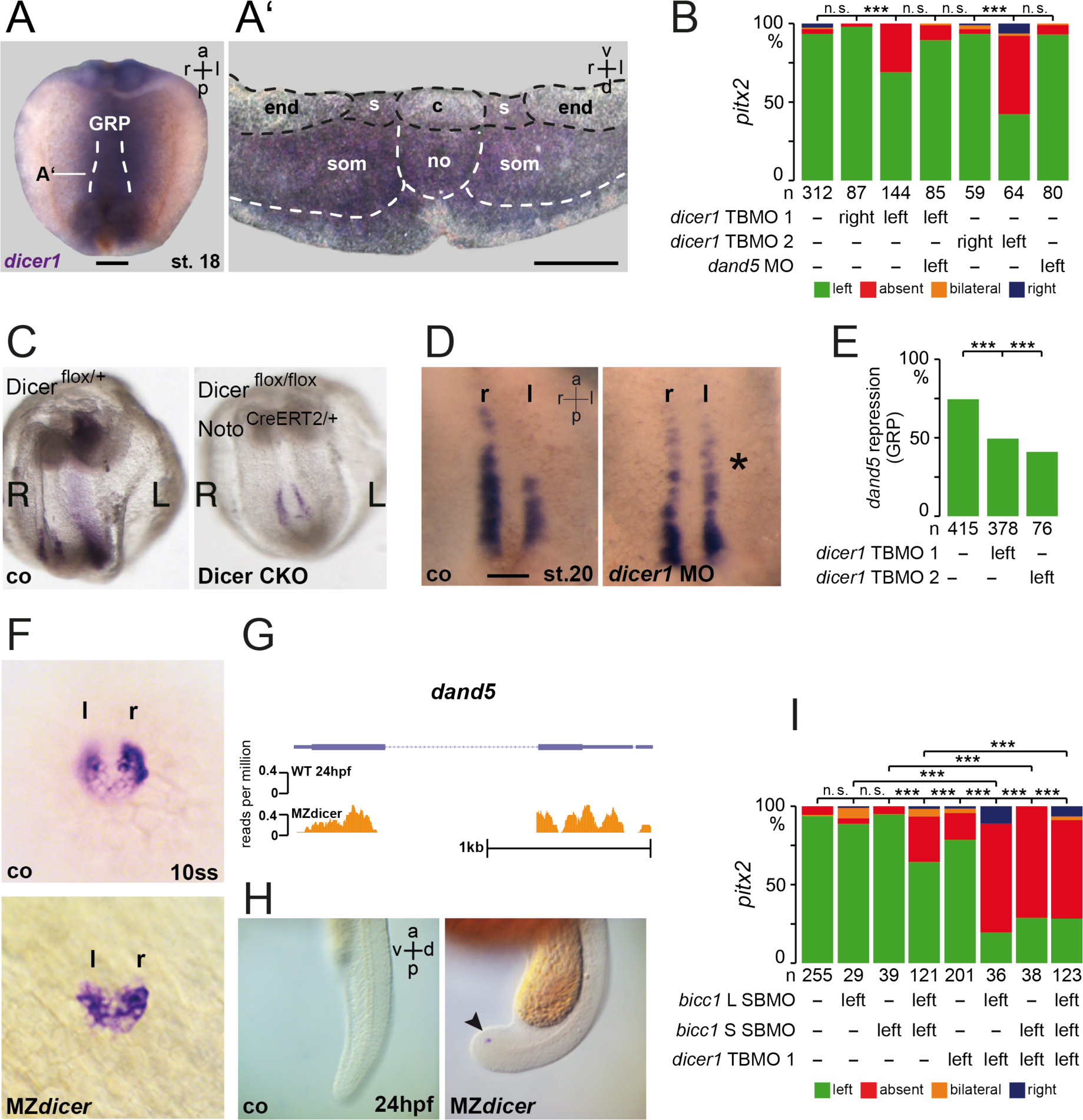
Dicer interacts with Bicc1 in *dand5* repression. (A) Expression of *dicer1* in sensory cells of the frog LRO (GRP; gastrocoel roof plate). Whole-mount in situ hybridization of a stage (st.) 18 dorsal explant with a *dicer1*-specific antisense RNA probe. (A’) Transverse histological section (level indicated in A) reveals mRNA expression in sensory (s) GRP cells, somites (som) and deep cells of the notochord (no), but absence of signals from central (c) flow-generating GRP and lateral endodermal cells (end). (B) MO-mediated inhibition of *dicer* translation in left, but not right sensory cells of the GRP prevented *pitx2* expression in the left LPM, which was rescued by a parallel knockdown of *dand5*. (C) mRNA expression of *Nodal* in control *(Dicer^flox/+^)* and *dicer* conditional knockout *(Dicer^flox/flox^ Noto^CreERT2/+^)* embryos at E8.0. Note that left-sided expression of *Nodal* in the left LPM was lost in the conditional mutants. (D) Absence of *dand5* mRNA decay at the left GRP margin in post-flow (stage 20) *dicer1* morphants. Representative dorsal explants of WT (left) and *dicer1* morphant (right) specimens hybridized with a *dand5*-specific antisense RNA probe. *, absence of decay. (E) Quantification of results. (F) Lack of *dand5* repression in 10 somite MZ*dicer* mutant zebrafish embryos. (G) Absence of *dand5* mRNA by RNAseq reads in 24hpf WT zebrafish embryos, but maintenance in MZ*dicer* mutants. (H) Absence of *dand5* mRNA by *in situ* hybridization in 24hpf WT zebrafish embryos, but maintenance in MZ*dicer* mutants (arrowhead). (I) *bicc1* and *dicer1* interact in *dand5* repression. Attenuated repression upon low-dose isolated injection of *bicc1* MOs as well as *dicer1* MO switched to high-level repression when *bicc1* and *dicer1* MOs were combined. a, anterior; d, dorsal; l, left; r, right; v, ventral. Numbers in (B, E, I) represent analyzed specimens from >3 independent experiments. n.s., not significant; ***, very highly significant, p<0.001. Scale bars in (A, D) represent 100 µm.

Analyzing earlier stages of laterality determination, downregulation of *dand5* mRNA levels at post-flow stages was compromised in *Xenopus dicer1* morphants (Figure 4D, E). This finding was conserved in zebrafish. In WT 10-somite stage embryos, *dand5* was repressed on the left side of Kupffer’s vesicle (KV), while no repression was observed in maternal-zygotic *dicer* mutants (Figure 4F). In the absence of Dicer, *dand5* expression was retained as late as 24hpf, a time when *dand5* expression was absent in wildtype embryos, as shown by RNAseq (Figure 4G) and whole-mount *in situ* hybridization (Figure 4H). Loss of *dand5* repression upon *dicer* loss-of-function could be caused by absence of flow or represent a miR-specific function.

MiRs have been shown to control motile ciliogenesis ^41^. In agreement with previous reports, which demonstrated a role of miRs in ciliogenesis in *Xenopus* ^42, 43^, ciliation of multiciliated cells in the *Xenopus* epidermis was impaired in *dicer1* morphants (not shown). When *dicer1* MOs were targeted to flow-generating GRP cells (C1-lineage), ciliation was unaltered in morphants (Figure S3D-F), demonstrating that *dicer* acted downstream of flow and upstream of *dand5* repression, like *bicc1* (and *pkd2* in mouse; ^44^. Next, we investigated whether *dicer1* and *bicc1* acted in the same pathway in flow sensing cells (C2-lineage). *bicc1* SBMOs (targeting S- and L-alleles) and *dicer1* TBMO1 concentrations were lowered such that when injected separately the result was a comparably low impairment of *pitx2* induction in the left LPM (Figure 4I). Co-injection of either *bicc1* S- or L-SBMO and *dicer1* MO, however, erased *pitx2* expression in about 70% of cases (Figure 4I), demonstrating that *bicc1* and *dicer1* synergize to mediate *dand5* repression. These results indicate that *dicer* and thus miRs are involved in *bicc1*-dependent posttranscriptional repression of *dand5*.

Finally, we wondered whether *pkd2*, one of two published active components in the flow sensor ^16, 44^, acted in the same pathway as well. Our recent demonstration of an earlier (likely maternal) *Pkd2* function in the specification and morphogenesis of the LRO prevented us from investigating this question in the context of LR axis formation in the embryo itself ^45^. In zebrafish, however, zygotic *pkd2* mutant embryos have strongly delayed induction of *Nodal*, but show normal KV ciliation and morphology ^46, 47^, suggesting a role for TRPP2 in flow sensing. In agreement with this notion, *dand5* mRNA repression was not observed in *pkd2* mutant and morphant zebrafish embryos (Figure 5A, B), likely leading to the strong delay in *Nodal* induction observed in these backgrounds (Schottenfeld et al., 2007). To test a potential interplay between *pkd2* and *bicc1* in the process of *dand5* repression, we returned to the animal cap reporter assay in *Xenopus* (Figure 2A). In order to be able to record additive effects of *pkd2*, we attenuated the *bicc1*-mediated repression of the *dand5*-reporter by lowering the concentration of co-injected *bicc1* mRNA, such that reporter activity was only repressed to some 40% of WT (Figure 5C). Upon co-injection of full-length *pkd2* mRNA, reporter activity was further repressed to less than 20% (Figure 5C). Because *pkd2* is maternally expressed in animal tissue, like *dand5* ^45^, we tested this interaction further by co-injecting *pkd2* MO, the specificity of which we showed previously ^27, 45^. Loss of *pkd2* partially rescued *bicc1*-mediated repression of the Luciferase *dand5* reporter (Figure 5C), establishing a firm link between *bicc1*, *pkd2* and post-transcriptional regulation of *dand5*.

**Figure 5.**
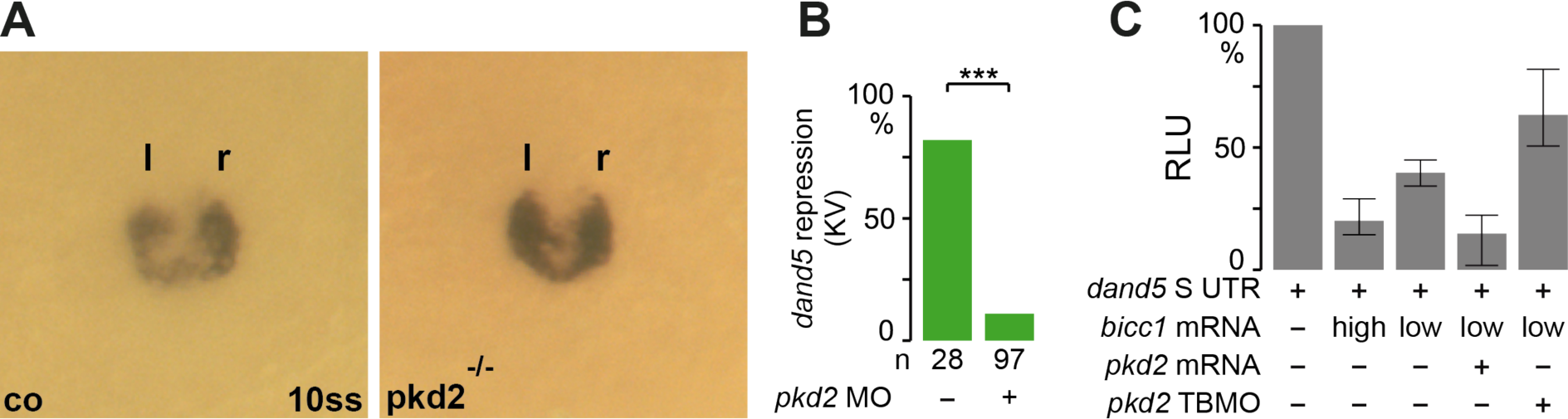
*Bicc1* and *pkd2* interact in translational repression of *dand5* (A) Absence of *dand5* repression in *pkd2* mutant zebrafish. (B) Absence of *dand5* repression in *pkd2* morphant zebrafish at 10ss. (C) Animal cap luciferase reporter assay of full-length *dand5* 3’-UTR (S-allele). The reporter construct was injected as mRNA into the animal region of 2-4 cell embryos, alone or in combination with high or low dose *bicc1* mRNA, *pkd2* mRNA or *pkd2* TBMO. Following culture to stage 10, animal cap tissue was excised and processed for determination of reporter protein activity (cf. Figure 2A). Attenuated repression upon co-injection of low concentrations of *bicc1* was reverted to high-level repression when *pkd2* mRNA was co-applied, or further diminished upon knockdown of *pkd2* using TBMO.

In summary, data presented here demonstrate that the RNA binding protein *bicc1* and the miR-processing enzyme *dicer* interact in flow-dependent *dand5* repression and cooperate with the calcium channel *pkd2* in sensing of leftward flow at the left LRO margin.

## DISCUSSION

LROs are quite peculiar structures which form and disappear in passing. They derive from superficial cells, function as LROs while embedded in the gut endoderm and – at least in the frog *Xenopus* – are destined to contribute to mesodermal tissues: notochord (medial flow generator) and somites (lateral flow sensor; ^2, 48^. They serve no other purpose than symmetry breaking ^49^ and may be needed as a flow-producing and -sensing tissue for no longer than ± 2 hours, as cells integrate fast into the forming somites and notochord with the endoderm closing above ^48^. Despite quite extensive variability in appearance, from the KV of bony fish to the GRP of amphibian and the ‘ventral node’ (posterior notochord) of mammalian embryos, LROs represent evolutionary conserved entities ^50^.

Many of the basic mechanisms of symmetry breaking at the ciliated vertebrate LRO have been solved through genetic and experimental studies in the various model organisms. The emerging general picture involves gastrula-stage pre-patterning of the LRO through organizer-derived mesodermal signaling cascades ^45, 51^; induction of motile ciliogenesis in the LRO center through activation of a transcriptional network including *foxj1* and *rfx* genes ^52^; polarization of cilia through Wnt/PCP-signaling ^53, 54^; and generation of leftward flow through rotating motile cilia; flow-mediated down-regulation of *dand5* on the left LRO margin as a prerequisite of signal transfer to the left LPM and induction of the Nodal signaling cascade ^1, 55^. A central remaining question awaiting a solution is the mechanism of flow-sensing resulting in *dand5* repression. While considerable effort is being invested in addressing how cilia may sense flow directionality (2-cilia vs. morphogen model; ^11, 56–59^, there is conflicting data regarding the involvement of mechano-sensory cilia in flow-sensing cells ^60–62^.

Therefore, we have chosen to tackle the problem starting from the downstream target process, namely the repression of *dand5*. Our work, together with complementing analyses in the mouse (cf. accompanying manuscript by Minegishi et al.), constitutes a conceptual advance in our understanding of symmetry breaking, namely the flow-dependent activation of the RNA-binding protein Bicc1 to repress *dand5* translation on the left LRO margin in a dicer-dependent manner. Based on our analyses, we like to suggest a model schematically depicted in Figure 6.

**Figure 6.**
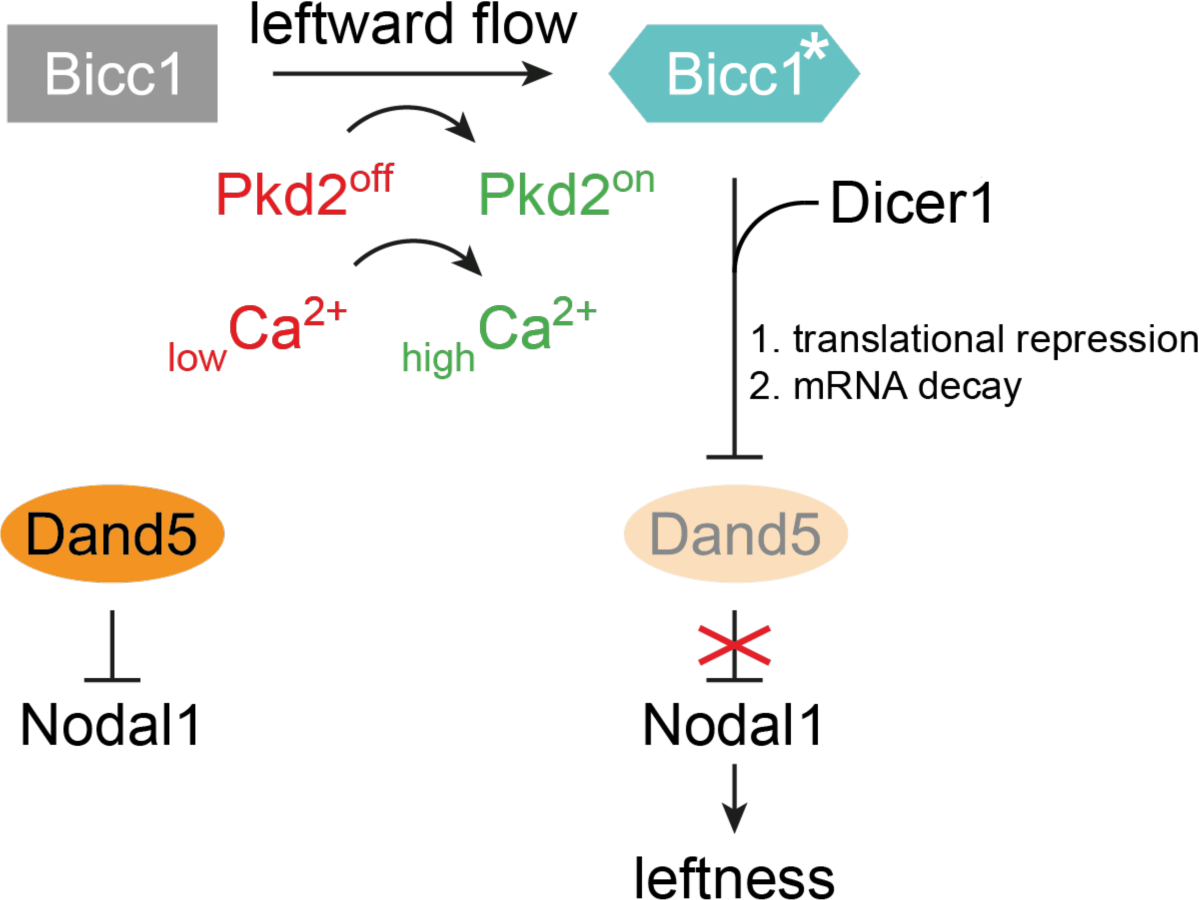
**Posttranscriptional repression of *dand5* in left flow sensor cells at the *Xenopus* left-right organizer: a model**. Leftward flow activates the TRP channel Pkd2 on the left side of the LRO, resulting in an asymmetric calcium signal in flow sensor cells. A calcium-dependent mechanism activates Bicc1 to become a repressor of *dand5* by dicer-dependent translational repression and mRNA decay. Attenuated Dand5 expression lifts repression of Nodal and defines leftness by induction of the LPM Nodal signaling cascade. For details, see text.

We hypothesize that prior to flow, Bicc1 does not interfere with *dand5* mRNA translation, a reasoning supported by the lack of a gain-of-function phenotype (Figure 1C and data not shown). Dand5 and Nodal are translated, with Dand5 repressing Nodal, probably requiring an excess of Dand5 protein. During flow stages, TRPP2 channel activation results in a cytoplasmic Ca^2+^ signal, which has been described in mouse and zebrafish ^16, 57, 62, 63^ and which represents the intracellular second messenger of the initially extracellular flow signal. We hypothesize that Bicc1 gets activated (Bicc1*) by TRPP2 and Ca^2+^, possibly by a Ca^2+^-dependent phosphorylation event to inhibit *dand5* mRNA translation during flow stages, followed by mRNA decay (Bicc1*; Figure 6). Bicc1 protein domains (cf Figure 2B) have been functionally characterized in various contexts, for example during *Drosophila* and *Xenopus* axis formation as well as mouse kidney development ^25^. KH and KH-like domains as well as the SAM can bind RNA; in addition, the SAM domain mediates protein-protein interactions, including self-polymerization ^31^. Down-regulation of *dand5* releases Nodal repression, which initiates LPM nodal cascade induction and asymmetric organ morphogenesis, i.e. leftness.

Bicc1 activation to Bicc1* should depend on *pkd2*, based on the described epistasis between *pkd2* and *bicc1* in *Xenopus* and the dependence of *dand5* repression on *pkd2* in zebrafish (cf. Figure 5). In *Drosophila*, Bicc1 interacts with and is phosphorylated by the Thr kinase complex PNG ^64, 65^. In the light of (a) these reports; (b) the *pkd2* dependence of Bicc1*; (c) predicted phosphorylation sites in Bicc1 (not shown), (d) and the responsiveness of *dand5* repression on calcium signaling in the mouse (cf. accompanying manuscript by Minegishi et al.), we envision that Ca^2+^ -dependent kinases phosphorylate Bicc1 to activate repression (Bicc1*) of *dand5*. In zebrafish, transient activation of CaMK-II downstream of asymmetric Ca^2+^ was shown to be required in the LRO (KV) for asymmetric Nodal cascade induction and correct development of organ situs ^66^.

Besides our experimental data, human genetics supports the notion that Bicc1 and TRPP2 cooperate in sensing leftward flow: mutations in both *pkd2* and *bicc1* give rise to autosomal dominant polycystic kidney disease in humans (ADPKD; ^23, 27^. As TRPP2 is considered to be the flow sensor in renal cells of the collecting ducts, we imagine that a *pkd2*/*bicc1* module acts in flow sensing during laterality specification as well. Flow-sensing at the vertebrate LRO represents an extremely dynamic process: cilia become motile and generate a leftward extracellular fluid flow for only a very short period of maybe 2-3 hours or even less, before the Nodal cascade gets asymmetrically induced in the left LPM and laterality becomes fixed. It is thus of central importance that processes that inactivate *dand5* act fast and direct, without the need of transcriptional activation of novel genes. The mechanism described here, involving Ca^2+^ as second messenger and phosphorylation of Bicc1 as a key target protein fulfil the required criterion of speed. This mechanism guarantees the efficient translational repression and mRNA degradation of *dand5* from LRO sensor cells.

In evolutionary terms, the *pkd2*/*bicc1*/*dicer* module is functionally conserved from zebrafish to mammals. In the mouse, like in *Xenopus*, a proximal element of the *dand5* 3’-UTR is required and sufficient for flow-mediated mRNA decay (mouse) and translational inhibition (frog), which is dependent on *bicc1* and *dicer* (cf. accompanying manuscript by Minegishi et al.). Whether or not microRNAs are involved in Dicer1-mediated *dand5* repression remains an open question. Down-regulation of *miR-15a*, a known regulator of Nodal signaling ^67^, has been linked to cyst formation in polycystic liver disease patients ^68^, which – from a human genetics point of view – seems to support a possible involvement of miRs. The generally long *dand5* 3’-UTRs, however, lack significant evolutionary sequence conservation between – and even within – the different classes of the vertebrates, at least in amphibians, where *X. tropicalis* differs greatly from *X. laevis*. Interestingly, this low conservation extends to the amino acid sequence of Dand5 proteins, which is rather low (<35% between human and *Xenopus*, human and zebrafish as well as *Xenopus* and zebrafish; about 60% between human and mouse). In the light of the functional conservation during symmetry breaking this is a surprising finding. What differs between the vertebrates that use cilia for symmetry breaking as well is – despite its functional conservation – the anatomy of LROs: the zebrafish KV presents as a closed sphere while in medaka the KV has a dome-like shape, amphibian GRPs are embedded in the archenteron and mammalian LROs, i.e. ventral nodes, are continuous with the posterior notochord ^69^. Transient LRO cells may differ in fate as well, which has not been addressed in great detail except for amphibians, where flow-sensing lateral GRP cells are of somitic fate ^48^. However, even in *Xenopus*, flow sensing cells differ in the way they integrate into the somites between *X. laevis* and *X. tropicalis* (ingression vs. relamination; ^48^.

Using our target protector MO, we were able to separate *dand5* mRNA decay from translation inhibition: left-sided, flow dependent *dand5* mRNA repression was still observed in tpMO-injected specimens (Figure 3C), while left nodal cascade induction was inhibited (Figure 3B. This result strongly suggests that a discrete regulatory mechanism of *dand5* mRNA stability is at work. Interestingly, binding sites for Bicc1 in the critical region of mouse *dand5* 3’-UTR, identified in the accompanying study (Minegishi et al.), are located within (*dand5-*S) or next to (*dand5*-L) the target protector MO sequences that impair repression in *Xenopus* (Figure S2C). Analyzing the proximal regions of the various *dand5* 3’-UTR sequences, which show the highest degree of conservation (cf. Figure S2), using different online miR-target prediction tools, only very few potential miR-binding sites show up, the probability of which is low in every single case. The one that may be of some significance is miR-133, because all members of this family are specific for muscle development and expressed in somites ^70–72^. A conserved target site was detected in *X. laevis* S- and L-alleles as well as in the human proximal *dand5* 3’-UTR (Figure S2C). It remains to be seen whether one of the four family members in *Xenopus* is involved in Bicc1-mediated *dand5* repression in *Xenopus*, where flow-sensing LRO cells are of somitic fate, which is unamenable to determine in humans.

Interestingly, it has been shown that Bicc1 regulates its own expression in a post-transcriptional manner ^65^. A highly conserved miR-133 binding site in 3’-UTRs of vertebrate *bicc1* genes (not shown) may suggest that a Bicc1/miR-133 module has been adapted to the regulation of *dand5* in somitic/flow-sensing LRO cells. Alternatively, Dicer1 may act miR-independently through one of its described non-canonical mechanisms ^73^.

In conclusion, our work identifies Bicaudal C and Dicer as two novel factors in sensing of leftward flow. They do not impact on specification of the LRO or flow generation, however their loss can be compensated by knockdown of *dand5*, the central process catalyzed by flow. The exact nature of Bicc1’s interaction with the *dand5* 3’-UTR remains to be solved. The evolutionary conservation of the molecular sensor module as well as its link to the ciliary calcium channel TRPP2, however, establish a firm link between extracellular leftward fluid flow and molecular symmetry breaking – *dand5* repression – within flow sensor cells.

Therefore, whatever flow brings about, ciliary bending or trafficking of cargo-laden vesicles (or both), should ultimately converge on TRPP2 activation.

## METHODS

### Plasmid construction

The m*bicc1*-CS2+ construct was a gift from Oliver Wessely (Cleveland, OH, United States). For *in vitro* synthesis of mRNA using the Ambion sp6 message kit, the plasmid was linearized with NotI. For *in vitro* synthesis of the luciferase reporter mRNA using the Ambion T7 message kit, the plasmid was linearized with BamH1.

### MO sequences and dosages of injections

*dand5-TBMO (Coco1-MO;* ^74^: 0.5 pmol per embryo

5’ – CTGGTGGCCTGGAACAACAGCATGT – 3’

*Dicer1-TBMO1* ^75^: 1.5 pmol per embryo

5′-TGCAGGGCTTTCATAAATCCAGTGA-3

*Dicer1-TBMO2* L covers ATG: 1 pmol per embryo

5′-CATGAGCTGAAGTCCTGCCATGC-3

*bicc1-SPMO1* L covers splice donor site exon 2: 1 pmol per embryo, low 0.5 pmol per embryo

5′-GGGAATAGACTCACCCTGTAACATT-3

*bicc1-SPMO2* (S) covers splice donor site exon 2: 1 pmol per embryo, low 0.5 pmol per embryo

5′-CCCAACAAGCAAGCTCTTACCTTCT-3

*bicc1-TBMO1* S (xBic-C-MO1; ^29^: 1 pmol per embryo

5′-TAG ACT CGC ACT GAG CCG CCA TTC T-3′ bicc1 TBMO L (xBic-C-MO2;^29^: 1 pmol per embryo

5′-CCA TTG TGC TAC TGC CGC CGC TAA C-3′

Pkd2 TBMO ^27^: 1 pmol per embryo

5′-GCCACTATCTCTTCAATCATCTCCG-3′

zfPkd2 TBMO ^46^: 1-4ng per embryo

5′-AGGACGAACGCGACTGGAGCTCATC-3′

dand5 tpMO S 3’-UTR: 1 pmol per embryo

5′-3. AAGTCGTCAAGTCGTTGGCACTTCC-3′

dand5 tpMO L 3’-UTR: 1 pmol per embryo

5′-TAGCACTTCCCCTGCTTCAGCAAAG-3′

### *Xenopus* frogs and embryos

Animals were handled in accordance with German regulations (Tierschutzgesetz) and approved by the Regional Council Stuttgart (A379/12 Zo, ‘Molekulare Embryologie’, V340/17 ZO and V349/18 ZO, ‘*Xenopus* Embryonen in der Forschung’).

*Xenopus* embryos obtained by *in vitro* fertilization were maintained in 0.1X modified Barth medium ^76^ and staged according to ^77^. During injections, embryos were kept in 1 x modified Barth medium with 2% Ficoll. To specifically target the sensing cells of the GRP for all experiments except for the luciferase assay, we injected into the dorsal marginal side (left or right; C2 lineage). For luciferase assays, embryos were injected twice into the animal blastomeres at the 4-cell stage with a luciferase *dand5* 3’-UTR construct, alone or together with a *bicc1* construct. Animal cap tissue was dissected at stage 10 (cf. Figure 2A for a schematic depiction of the procedure). Following injections, all embryos were transferred to 0.1 modified Barth medium.

### Zebrafish

Established husbandry protocols where adhered to, and experimental protocols conducted, in accordance with the Princeton University Institutional Animal Care and Use Committee (IACUC) guidelines. Embryos were raised at 28°C and processed for injections and RNA *in situ* hybridization as described ^46^. Zebrafish strains utilized include *pkd2/cup^tc^*^321^ ^46^ and *dicer1^hu^*^715^ ^78^. Embryos were staged according to ^79^.

### Immunfluorescence staining

For immunofluorescence staining, embryos were fixed in 4% PFA for 1h at RT on a rocking platform, followed by 2 washes in 1x PBS^-^ for 15 min each. For staining of GRP explants, embryos were dissected using a scalpel into anterior and posterior halves. Posterior halves (GRP explants) were collected and transferred to a 24-well plate and washed twice for 15 min in PBST. GRP explants and whole embryos were blocked for 2h at RT in CAS-Block diluted 1:10 in PBST. The blocking reagent was replaced by antibody solution (anti-acetylated tubulin antibody, diluted 1:700 in CAS-Block) and incubated over night at 4°C. In the morning, the antibody solution was removed and explants/embryos were washed twice for 15 min in PBS^-^. The secondary antibody (diluted 1:1000 in CAS-Block) was added together with Phalloidin (1:200) and incubated for a minimum of 3hrs at RT. Before photo documentation, embryos or explants were briefly washed in PBS^-^ and transferred onto a microscope slide.

### Luciferase assay

Luciferase reporter assays were carried out using the Promega Dual-Luciferase® Reporter Assay System. Animal cap tissue was transferred into a 1.5 ml Eppendorf tube and the 0.1xMBSH buffer was removed, leaving the tissue moistened. The tissue was lysed and homogenized in 100 µl 1X passive lysis-buffer by pipetting the suspension up and down, followed by 15 min incubation at RT. The lysate was centrifuged for 2 minutes at 14 000 rpm and the upper phase was transferred into a new tube. The lysate was re-centrifuged and two 25 µl aliquots (technical duplicates) of each sample were transferred into a 96-well plate. 75 µl 1x Luciferase assay substrate was added through the GloMax® Explorer System and luminescence was determined. This step was repeated with 75 µl 1X Stop and Glow reagents.

To calculate the relative luciferase units (RLU [%]), the ratio between luciferase and Renilla values was calculated and correlated to the WT control, which was set to 100%.

### Statistical analysis

Statistical calculations of marker gene expression patterns and cilia distribution were performed using Pearson’s chi-square test (Bonferroni corrected) in statistical R. For the statistical calculation of ciliation, a Wilcoxon-Match-Pair test was used (RStudio).

### Mouse Strains

All mouse experiments were performed in accordance with guidelines of the RIKEN Center for Biosystems Dynamics Research (BDR) and under an institutional license (A2016-01-6). Mice were maintained in the animal facility of the RIKEN Center for BDR. *Noto-Cre*^ERT2^ mice were described in ^80^, *Dicer^flox^* mice in ^81^, JAX stock #006001). Expression of the *Noto-Cre*^ERT2^ transgene in embryos was induced by oral administration of tamoxifen (Sigma) in corn oil to pregnant mice at a dose of 5 mg both 24 and 12 h before the late headfold stage.

### WISH Analysis in mouse

WISH was performed according to standard procedures with digoxigenin-labeled riboprobes specific for *Nodal* mRNA ^82^.

### DATA AVAILABILITY

The authors declare that the main data supporting the findings of this study are available within the article and its Supplementary Information files.

## ACKNOWLEDGEMENTS

We thank M. Montino for performing initial Bicc1 experiments in *Xenopus* during his diploma thesis. O. Wessely provided the full-length mouse *bicc1* construct. We thank H. Sasaki for *Noto*^CreERT2^ mice and Philip Johnson for zebrafish husbandry. MG was the recipient of a Ph.D. fellowship from the Landesgraduiertenförderung Baden-Württemberg. Work in the Blum lab was supported by DFG grant BL285/9-2. Work in the Sheets lab was supported by the National Institute of Child Health and Human Development of the National Institutes of Health under award number R01HD091921. Work in the Hamada lab was supported by grants from the Ministry of Education, Culture, Sports, Science, and Technology (MEXT) of Japan (no. 17H01435) and from Core Research for Evolutional Science and Technology (CREST) of the Japan Science and Technology Agency (no. JPMJCR13W5) to H.H.; by a Grant-in-Aid (no. 18K14725) for Early-Career Scientists from the Japan Society for the Promotion of Science (JSPS), a Kakehashi grant from BDR-Otsuka Pharmaceutical Collaboration Center, and a research grant (no. 2018M-018) from the Kato Memorial Bioscience Foundation to K.M. Work in the Giraldez lab was supported by the National Institute of General Medical Sciences of the National Institutes of Health under award number R35GM12258003. Work in the Burdine lab was supported by the National Institute of General Medical Sciences of the National Institutes of Health under award number R01HD048584, and a United Negro College Fund/Merck Graduate Science Research Dissertation Fellowship to JCM. The content is solely the responsibility of the authors and does not necessarily represent the official views of the National Institutes of Health.

## AUTHOR CONTRIBUTIONS

Experiments in *Xenopus* were performed by MM (bicc1, dicer), MG (dicer), MD (part of AC assays) and PV (initial bicc1 work). Zebrafish experiments were conducted by JLP and JCM, RNAseq by VY. The conditional *dicer* knockout mouse was generated and analyzed by KM. AS, MB, HH, RDB, AJG and MS conceptualized and supervised experiments, which were analyzed by all authors. MB wrote the manuscript with suggestions from all authors.

## SUPPLEMENTAL FIGURES AND LEGENDS

**Figure S1.**
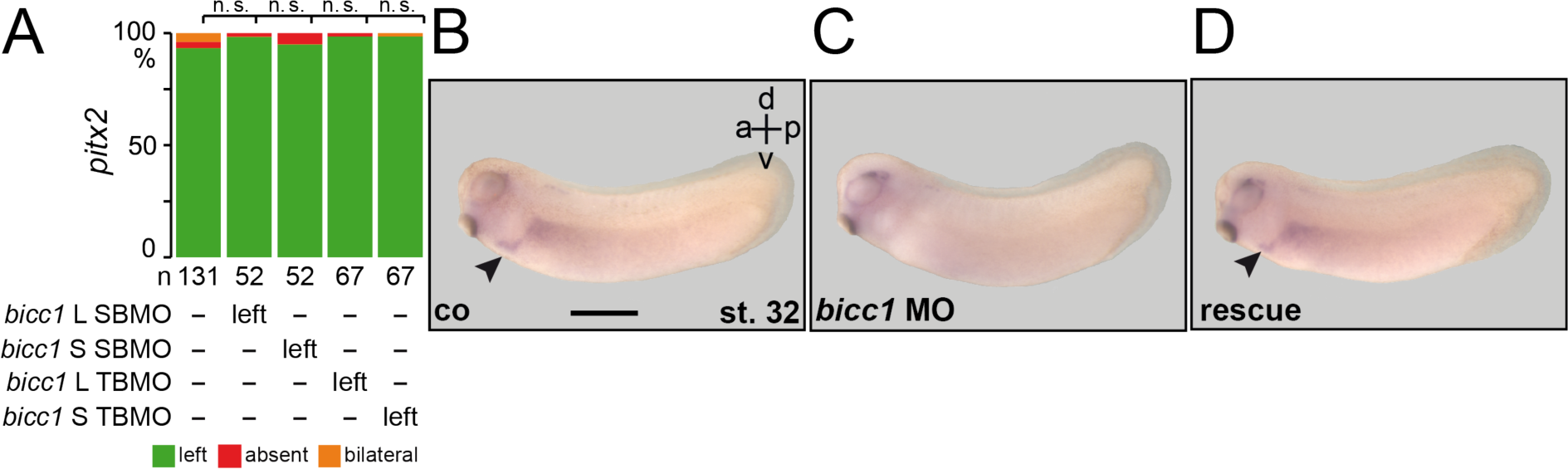
Characterization of *bicc1* morphants (A) Individual injection of L- or S-alloallele-specific SBMOs did not affect *pitx2* expression in morphants. (B-D) *pitx2* expression in representative control (co; B), *bicc1* morphant (C) and specimen in which SBMO and a full-length *bicc1*mRNA not targeted by the MO were co-injected (D). n.s., not significant. Scale bar in (B) represents 1mm.

**Figure S2.**
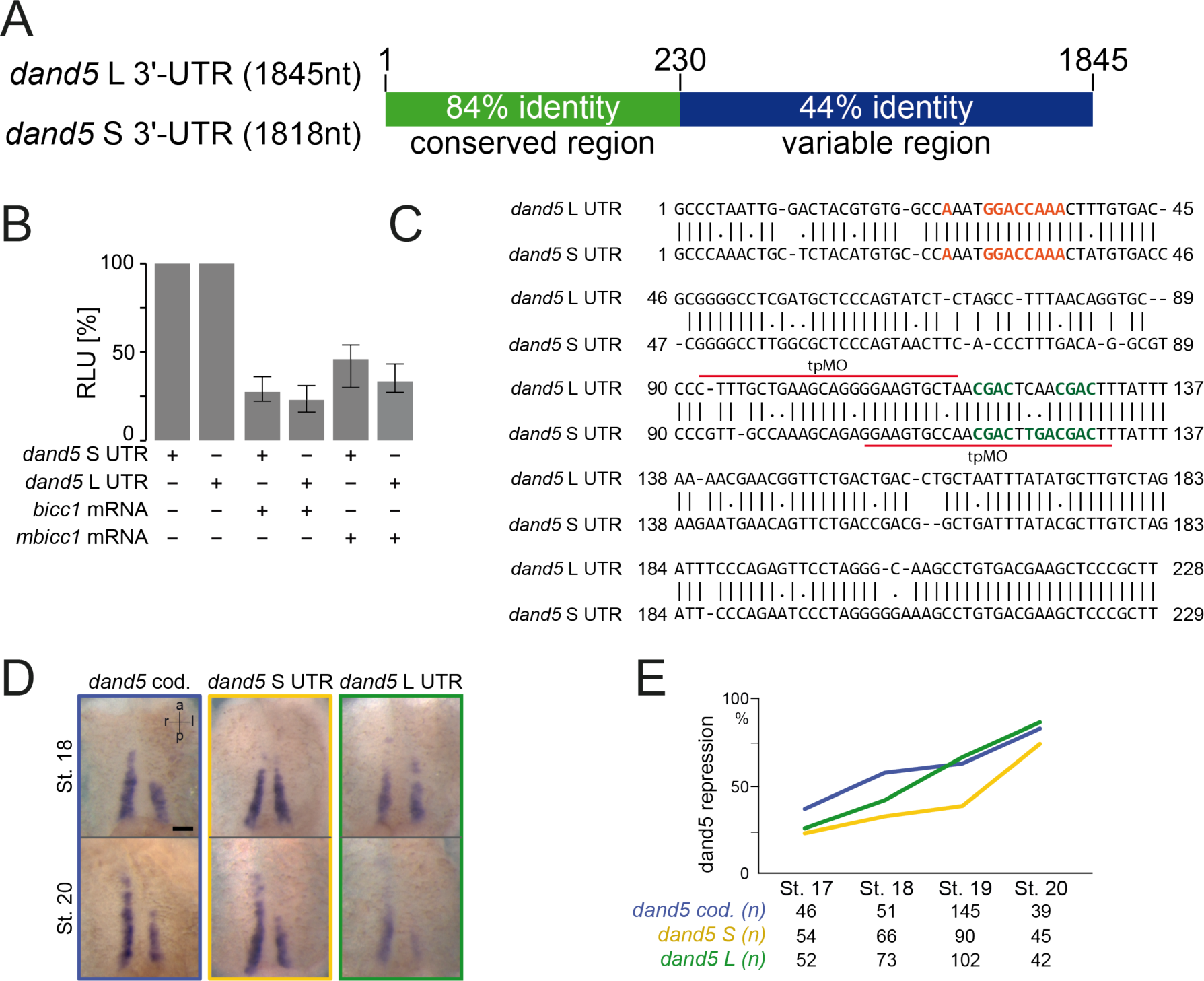
Characterization of *dand5* 3’-UTRs (A) Conservation of 3’-UTR sequences between S- and L-alloalles of *X. laevis*. (B) Animal cap reporter assay (cf. Figure 2A) following injections of *dand5* S- or L-3’-UTRs alone or together with *Xenopus* (*bicc1*) or mouse *bicc1* (*mbicc1*) effector mRNAs. Note that both alloalleles were equally repressed. Note also that *mbicc1*was efficient as a repressor as well. (C) Sequence alignment of the proximal 229 respective 228 nucleotides of *dand5* 3’-UTRs of S- and L-alloalleles. The position of the TPMOs used in experiments depicted in Figure 3B and 3C are marked by red lines+. (D) Representative dorsal explants of stage 18 (top row) and stage 20 (bottom row) embryos hybridized with antisense RNA probes specific for the *dand5* coding sequence (left), or the 3’-UTRs of *dand5* S-(middle) and L-allele (right). (E) Quantification of results of a time course analysis from stage 17-20. The graph displays left-sided repression of *dand5* expression following visual judgement of dorsal explants following *in situ* hybridization. Scale bar in (D) represents 100 µm.

**Figure S3.**
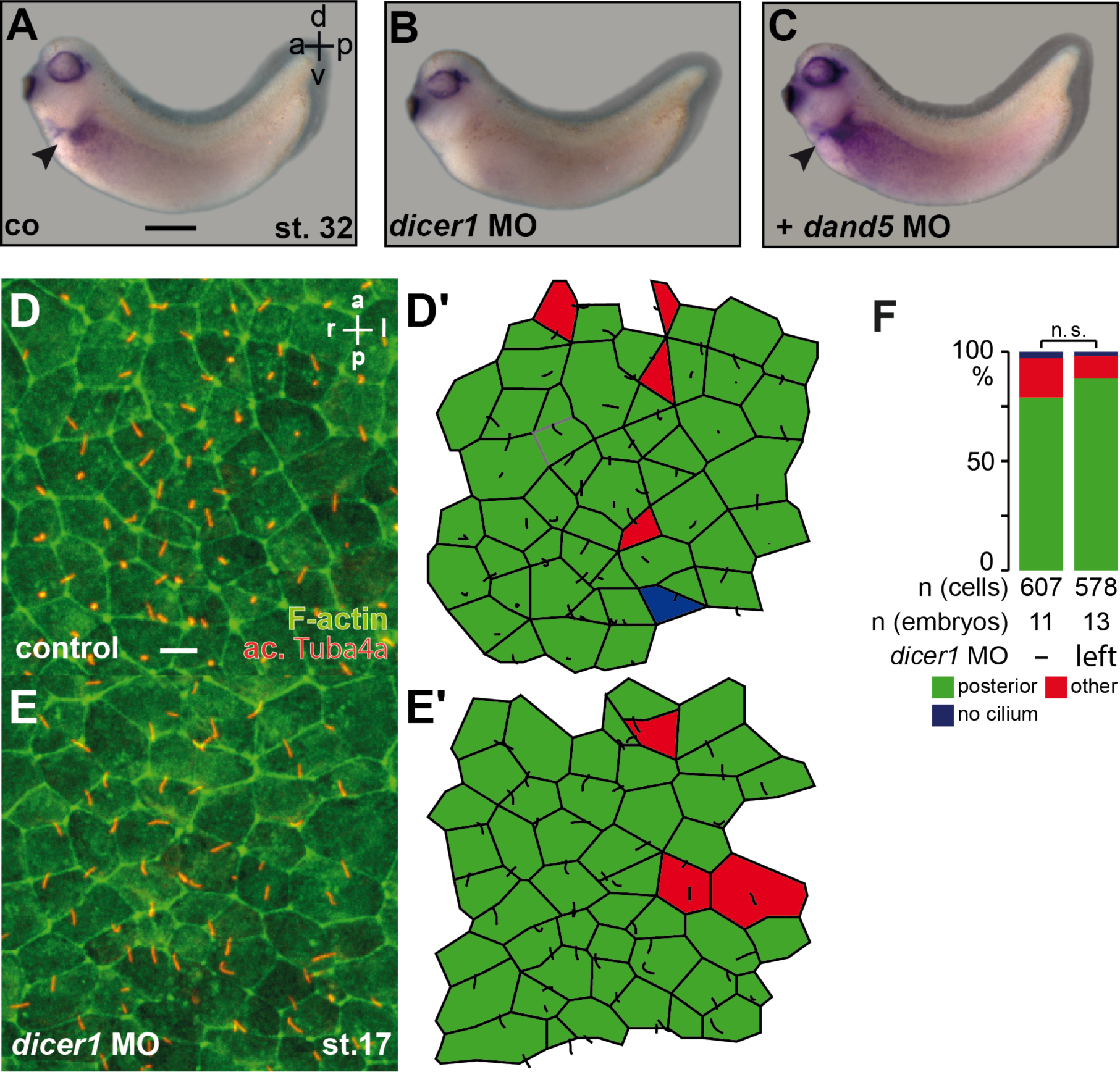
A post-flow role of *dicer1* in *Xenopus* LR axis formation. (A-C) Repression of *pitx2* expression in the left LPM of stage (st.) 32 *dicer1* morphant tadpoles was rescued by parallel knockdown of *dand5* translation. Whole-mount in situ hybridization of specimens with a *pitx2c*-specific antisense RNA probe. (D-F) WT morphology and ciliation of GRPs in *dicer1* morphant (E) as compared to uninjected control specimen (D), as shown by immunofluorescence staining of cilia using an antibody against acetylated tubulin (ac. Tuba4a; red) and counterstaining of actin using phalloidin to highlight cell boundaries (green). (D, E) Evaluation of cell morphologies and ciliation in dorsal explants. (F) Quantification of cilia polarization. Scale bars in (A) represents 1 mm. in (D) 10 µm.

